# The trade-offs of sharing pollinators: pollination service is determined by the community context

**DOI:** 10.1101/865279

**Authors:** E. Fernando Cagua, Hugo J. Marrero, Jason M. Tylianakis, Daniel B. Stouffer

## Abstract

A fundamental feature of pollination systems is the indirect facilitation and competition that arises when plants species share pollinators. When plants share pollinators, the pollination service can be influenced. This depends not only on how many partners plant species share, but also by multiple intertwined factors like the plant species’ abundance, visitation, or traits. These factors inherently operate at the community level. However, most of our understanding of how these factors may affect the pollination service is based on systems of up to a handful of species. By examining comprehensive empirical data in eleven natural communities, we show here that the pollination service is—surprisingly—only partially influenced by the number of shared pollinators. Instead, the factors that most influence the pollination service (abundance and visit effectiveness) also introduce a trade-off between the absolute amount of conspecific pollen received and the amount relative to heterospecific pollen. Importantly, the ways plants appear to balance these trade-offs depend strongly on the community context, as most species showed flexibility in the strategy they used to cope with competition for pollination.

## Introduction

Animal pollination plays a disproportionally important role in food production and maintenance of global biodiversity (Klein et al. 2007, Bascompte and Jordano 2007, Ollerton et al. 2011). At a pairwise level, the mutually beneficial relationship between plants and pollinators underpins the pollination service. At a community level, sometimes involving hundreds of species, both plant and pollinator species are connected in a myriad of indirect connections when pollination partners are shared. These indirect connections can dramatically alter the quality of the pollination service that plants receive because they determine how conspecific and heterospecific pollen is transferred across the community (Morales and Traveset 2008). Generally speaking, there is a trade-off between the benefits gained from a species maximising its number of partners and the costs of sharing them with other plant species (Waser 1978). However, due to the large number of factors that operate at the community level, we generally do not know how sharing pollinators affects the pollination service beyond systems with more than a handful of species. Here we investigate how pollinator sharing affects pollen transfer *in natural communities* and how it compares to other factors known to play a role in community dynamics like abundance, traits, and visitation patterns.

There are two main mechanisms through which sharing pollinators can affect plant fertilisation (Morales and Traveset 2008). The first is by changes in intraspecific pollen transfer. Changes in intraspecific pollen transfer happen, for example, when plants with more attractive flowers might reduce the number of visits to those less attractive neighbouring plants, and hence reduce the amount of *conspecific pollen* deposited by animals (Yang et al. 2011). The second is via interspecific pollen transfer. In that case, even receiving a visit might not necessarily translate into fertilisation (Campbell and Motten 1985) because a focal plant might receive *heterospecific pollen* or because pollen from the focal plant might be lost to different species. Naturally, the precise effects on female or male plant fitness of conspecific and heterospecific pollen deposition depend on the species involved (Arceo-Gómez and Ashman 2016) and are unknown for many plant species.

Even for species well adapted to pollinator sharing, receiving foreign pollen on stigmas or losing pollen to foreign stigmas is neutral (at best). Indeed, there is substantial evidence supporting the idea that heterospecific pollen deposition can be detrimental to seed production and plant fitness (Ashman and Arceo-Gómez 2013, Arceo-Gómez and Ashman 2016). All else being equal, provided pollen is viable and compatible (de Jong et al. 1992, Dafni and Firmage 2000, Ramsey and Vaughton 2000), the higher the quantity of conspecific pollen and its purity (relative to heterospecific pollen), the better the pollination service received by the focal plant. As such, measuring conspecific and heterospecific pollen deposition provides a good indication of the potential levels of facilitation and competition a plant population might experience.

By definition, intra- and interspecific pollen transfer occur at the community scale. However, with few exceptions (Aizen and Rovere 2010, Tur et al. 2016), most of what we know about pollen transfer and its relationship with key ecological factors are based on studies with two plant species. That is partly so because the factors that determine the patterns of pollen deposition at the community scale are tightly intertwined, operate simultaneously, and may lead to emergent phenomena not observed at smaller scales (Flanagan et al. 2011). For instance, recent empirical evidence suggests that plants with flowering traits that are “original” relative to others in the community generally have fewer interaction partners (Coux et al. 2016).

This evidence aligns with the notion that a species that interacts with few species does so strongly with each of them whereas a species that interacts with a large number of species does so comparatively weakly (Bascompte et al. 2006, Vázquez et al. 2007, Thébault and Fontaine 2008). If evolutionary specialisation occurs by changing traits to focus on fewer but better partners (Caruso 2000), we should expect a reduction of competition for pollinators in species with “original” traits and an increase of competition in species with a large number of interaction partners (Gibson et al. 2012, Carvalheiro et al. 2014). Alternatively, it might also be the case that abundance (for example, in terms of flower or pollen counts) is the dominant force driving pollen transfer (Seifan et al. 2014). Abundant plant species might experience a dilution of available pollinators (Feinsinger 1987, Feldman et al. 2004) but might also receive more effective visits by capitalising on a larger share of both visits and the pollen carried by pollinators (Stavert et al. 2019). In this case, a potential reduction in the absolute amount of conspecific pollen received could be compensated by an increase in the amount of conspecific pollen relative to heterospecific pollen. Altogether, it is clear that these ecological factors can indeed shape pollen deposition at the community level. However, we still do not understand their relative importance and *the trade-offs* that might exist between them.

Here, we investigate pollen-deposition dynamics at the community scale using empirical data from eleven plant-pollinator communities in the Argentinian Pampas. First, we investigate the relative contribution that four ecological factors make to the pollination service. Specifically, we hypothesise that there are trade-offs on how these factors affect the quantity and purity of conspecific pollen deposition. While quantity and purity should decrease for plants that share many pollination partners, other factors like the plant’s functional originality, its relative floral abundance, and its visitation patterns should have the potential to compensate for this decrease Second, we examine how these four factors that might affect pollen deposition can change across communities where species are present. Because these factors may affect the pollination service in contrasting ways, and a species role is relative to other species in the community, we predict that species present in multiple communities should be flexible enough to compete for pollinators under different community contexts.

## Methods

We collected data from eleven co-flowering plant communities and their pollinators in three locations in the Argentinian Pampas. In each location, we sampled two restored and two agricultural fragments, except in one located in the Flooding Pampas, where we were only able to sample one restored fragment due to the lack of available sites.

### Factors affecting quantity and purity of pollination service

Our first objective was to investigate the relative contribution that different ecological factors have on pollen deposition. Generally speaking, we expect that any factor that increases the amount of conspecific pollen deposited in stigmas, both in quantity and purity relative to heterospecific pollen, also has a positive effect on the pollination service. Specifically, we investigated the effect of (*i*) a plant’s number of shared pollinator species, (*ii*) a plant’s abundance relative to the rest of the community, (*iii*) the mean visit potential—a metric that combines the amount and type of pollen carried by floral visitors and the number of visits it receives from them, and (*iv*) the plant’s functional originality (Laliberté and Legendre 2010). See *Data Analysis* section below for more details on these four factors.

#### Data collection

In each of the studied communities, we quantified pollen deposition in a subset of plant species between December 2010 and February 2011. This subset comprised between three and nine common insect-pollinated (entomophilous) plant species that were flowering during the sampling period. Based on data from previous years (Marrero et al. 2014), we chose plant species such that they cover a wide range on a specialization-generalization gradient as well as a wide range of abundances. In each of the selected plants, we removed all flowers leaving only buds that were expected to go into florescence on the next day. Two days after flowering, we collected all remaining flowers and counted the number of conspecific and heterospecific pollen grains in their pistils. More details can be found in Marrero et al. (2016).

To obtain the number of shared pollinators for each species, we collected data to construct qualitative and quantitative pollination networks. Qualitative networks were constructed based on ten-hour observations of floral visits in each fragment. Quantitative networks were constructed using two 50 m randomly located transects in each fragment. We counted and collected all floral visitors found in a 2 m wide strip while walking at a pace of 10 m per minute (Memmott 1999, Marrero et al. 2014). We visited the transects each month between November 2010 and March 2011. To obtain floral abundance, we counted all units of floral attraction found during an independent sampling of the same transects used to construct the quantitative visitation networks. To estimate visit potential, we need to construct pollen transfer networks in addition to the visitation networks. To do this, we examined the pollen loads present on the floral visitors collected (Marrero et al. 2017). When the pollen count on an individual animal was estimated to be less than 2,000 grains, we identified every grain to the species level when possible and to pollen complexes when it was not. When the pollen count was above 2,000 grains, we classified approximately 50% of pollen and total pollen counts were extrapolated (Bosch et al. 2009). Finally, we also recorded morphological traits that relate to plant type (herb, shrub, climber), life cycle (annual, perennial), flower colouration, phenology, and whether the species is native in the study region. More details can be found in Marrero *et al.* (2014 and 2017).

#### Data analysis

To investigate the impact of ecological factors on pollination services, we used two sets of linear mixed models (LMM) with bootstrap resampling. The response variables for these model sets were the number of conspecific and heterospecific pollen grains deposited per stigma in flowers open to animal-mediated pollination. We used LMMs in which pollen loads were log-transformed because these models offered a better fit than equivalent GLMMs with Poisson (or quasi-Poisson) error structure. Models were fitted using the R package `nlme 3.1-131` (Pinheiro et al. 2018).

Because the amount of deposited pollen can vary widely across species, and potentially also across communities, we evaluated two possible structures for the random effects: one that includes a random intercept for plant species, and one that treats measures from species across different communities independently. We selected the best random structure by comparing the median Akaike Information Criterion for small samples (AICc).

As fixed predictors in the models, we included the four ecological factors described above. Specifically, we calculated the number of shared pollinator species for each plant species by pooling data from the qualitative and quantitative pollination networks. To calculate the plants’ relative floral abundance in their community, we aggregated floral counts for each species. We then calculated the mean visit potential of pollinator species *i* to plant species *j* as

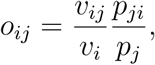

where *v*_*ij*_ is the observed number of visits by *i* to *j*, *p*_*ji*_ is the number of pollen grains from *j* attached to *i*, *v*_*i*_ is the total number of visits performed by *i*, and *p*_*j*_ is the total number of grains carried by *j*. We log-transformed the number of shared pollinators, floral abundance, and visit potential before including them in the model.

Finally, functional originality is defined as the distance of a species from the community trait average–the centroid of functional space of the community (Laliberté and Legendre 2010, Coux et al. 2016). To include phenological variation, we treated floral abundance in each of the survey months (November to March) as a “trait” in this analysis. To account for the non-independence of floral counts and weight all traits equally, we assigned a weight of 1/5 to these abundances (one for each month). We scaled all traits before calculating the centroid of the functional space and calculated the species-specific functional coordinates using the R package `FD 1.0-12` (Laliberté et al. 2014). Finally, we scaled all four factors to have a zero mean and unit variance.

To estimate the coefficients, perform model selection, and quantify the associated uncertainty, we used a combination of multi-model inference and bootstrap resampling with 99 replicates. Using bootstrap replicates allow us to better understand the uncertainties associated with our estimations. First, we performed model selection using AICc and determined the likelihood of each candidate model (a particular combination of predictors) by calculating the median ∆AICc (relative to the most likely model) for each bootstrap sample. As we wanted model coefficients from more likely candidate models to carry more weight in our results, we sampled the coefficients for our factors proportionally to the likelihood of their candidate model. Finally, we used these distributions of the model coefficients to estimate their mean impact on the pollination service (in terms of quantity and purity of conspecific pollen deposition).

### Flexibility of plant strategies

Our second objective was to tease apart whether and how these factors that might affect pollen deposition might change across communities species are present. If community context plays a relatively small role, or species are inflexible in regards to these factors, we would expect plants of the same species to use similar “strategies” across different communities. Alternatively, if the community plays a significant role and plant species are flexible, we should be able to observe differences in the strategy a plant species uses across communities. To test this, we first used a principal component analysis (PCA) of the four ecological factors (number of shared pollinators, floral abundance, visit potential, and trait originality). We scaled factors across the whole study to ensure that the PCA space does not change according to the species present in each community. We define a species’ strategy in a community as its coordinates in PCA space. For each species that was present in two or more communities, we then calculated (*i*) the median distance between the points that correspond to the strategy a species uses in different communities and (*ii*) the area of the convex hull defined by these points in the first two principal components (only for species present in three or more communities). We then compared the magnitude of these two metrics to those obtained with 99 Monte Carlo randomizations in which we replaced the strategy of the focal plant species by that of another randomly selected species in the dataset.

## Results

### Factors affecting quantity and purity of pollination service

We first examined the potential roles played in pollen deposition by four ecological factors (number of shared pollinators, abundance, mean visit potential, and functional originality). We found that our models of pollen deposition had high explanatory power (the coefficient of determination R^2^ ranged between 0.76 and 0.93) although a large portion of the explanatory power came from the random effects (Table S3). As determined by AICc, the random structure best supported by the data was the one that fit a separate intercept for each species in each community (as opposed to a common intercept for each species irrespective of the community to which they belong). This structure was best for both the models of conspecific and heterospecific pollen (Table S4).

Of the four factors we considered, we found that a plant’s mean visit potential and relative floral abundance were the most important at predicting pollen deposition in plant stigmas (Fig. 1a). Surprisingly, the number of shared pollinators was comparatively unimportant, particularly for models of heterospecific pollen deposition, as it was only ever included in models with relatively large AICc values (Table S5).

**Figure 1:**
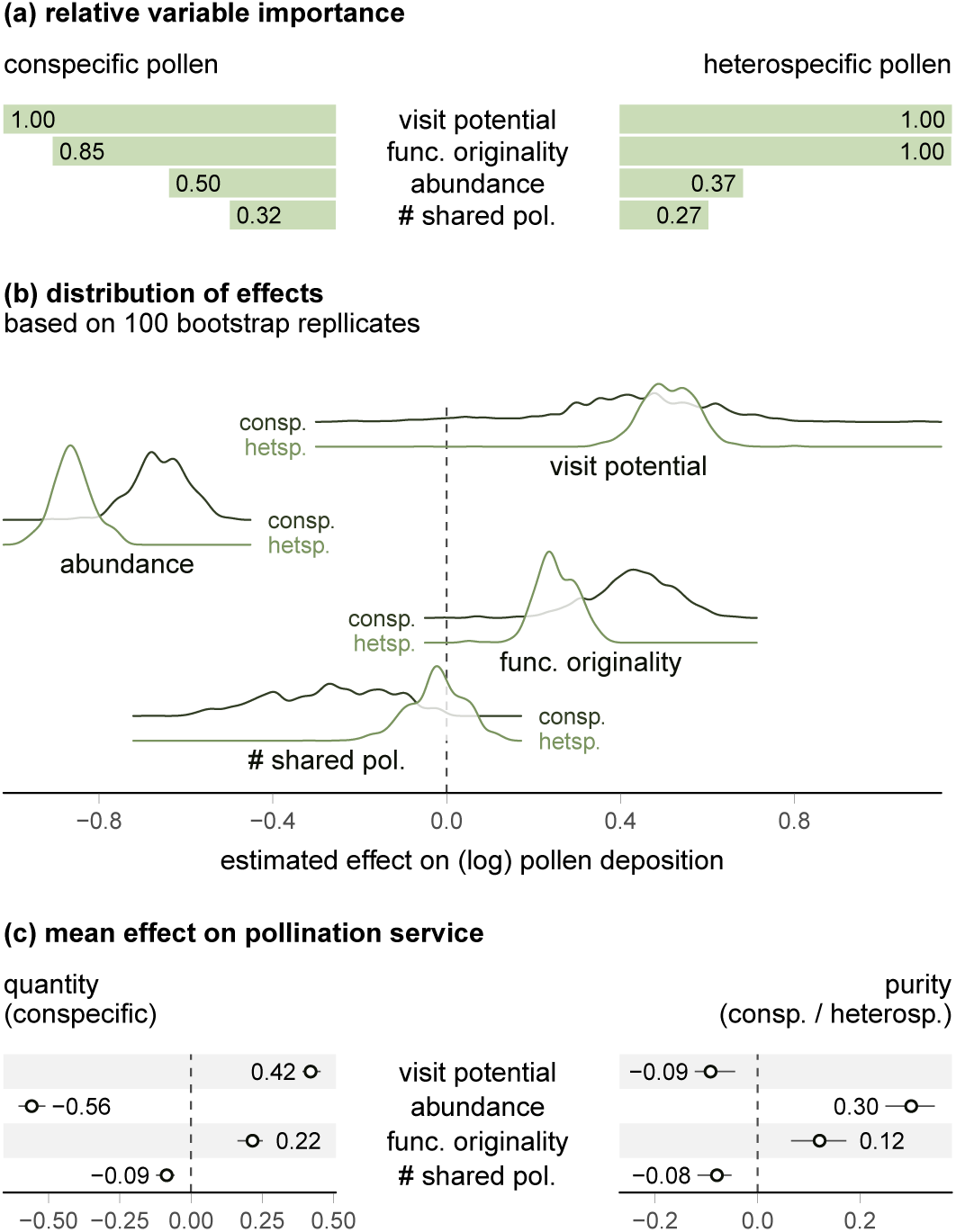
Effect of ecological factors on the pollination service. (a) The plant’s visit potential and relative floral abundance are the most important factors determining the deposition of conspecific and heterospecific pollen. Meanwhile, the number of shared pollinators was generally less important. The graph shows the relative importance calculated as the sum of the Akaike weights of the candidate models that included the selected factor. (b) The association between ecological factors and heterospecific pollen (lighter line) tended to align with their association with conspecific pollen (darker line). Visit potential and functional originality had a positive association with pollen deposition, while floral abundance and the number of shared pollinators had a negative association. The plot shows the distribution of the effects (across 99 bootstrap replicates) of the four ecological factors for conspecific and heterospecific pollen. (c) The end result of these associations is that only the plants’ functional originality has a positive impact on both the quantity and purity of conspecific pollen deposition (relative to heterospecific pollen). The plot shows the model averaged mean effect (*±* SE of 99 bootstrap replicates).

We found that the relationship between each of the ecological factors and pollen deposition was similar for both conspecific and heterospecific pollen. That is, strategies that were associated with an increase in conspecific pollen deposition were also associated with an increase in heterospecific pollen deposition. Specifically, the plants’ mean visit potential had a positive effect on pollen deposition (Fig. 1b). However, the effect size was slightly larger for heterospecific than for conspecific pollen. This larger effect indicates that, although there is a positive association between visit potential and the quantity of pollen deposition, there is a negative relationship with its purity (Fig. 1c). In contrast, a plants’ relative floral abundance negatively affected its deposition quantity, but the mean difference between the coefficients in the models indicates a positive association with purity (Fig. 1c). The third most important factor, functional originality, had a positive, although comparatively smaller, association with both the quantity and purity. Finally, the number of shared pollinators had negative and neutral associations with conspecific and heterospecific pollen, respectively, but these impacts were small when compared to the other factors. Although the ecological factors were positively correlated (Fig. S2), the collinearity between predictors did not qualitatively affect our findings (Fig. S3).

### Flexibility of plant strategies

We used a PCA of the ecological factors–species matrix to investigate whether plants’ strategies towards pollen deposition is similar across communities or whether they are flexible and therefore a reflection of the community context. The first two PCA components explained 75% of the total variance (Fig. 2a). The first component was dominated by visit potential and relative abundance while the second component was dominated by the number of shared pollinators and the plant’s functional originality. When we locate the species that were sampled in more than one community in the first two PCA components (Fig. 2b), we observe that the positions of any given species do not tend to be close to each other. Indeed, when we measured the median distance between the plants’ coordinates, we found that it was only significantly smaller than that of randomisations for only two of the twelve species analysed (Fig. 2c).

**Figure 2:**
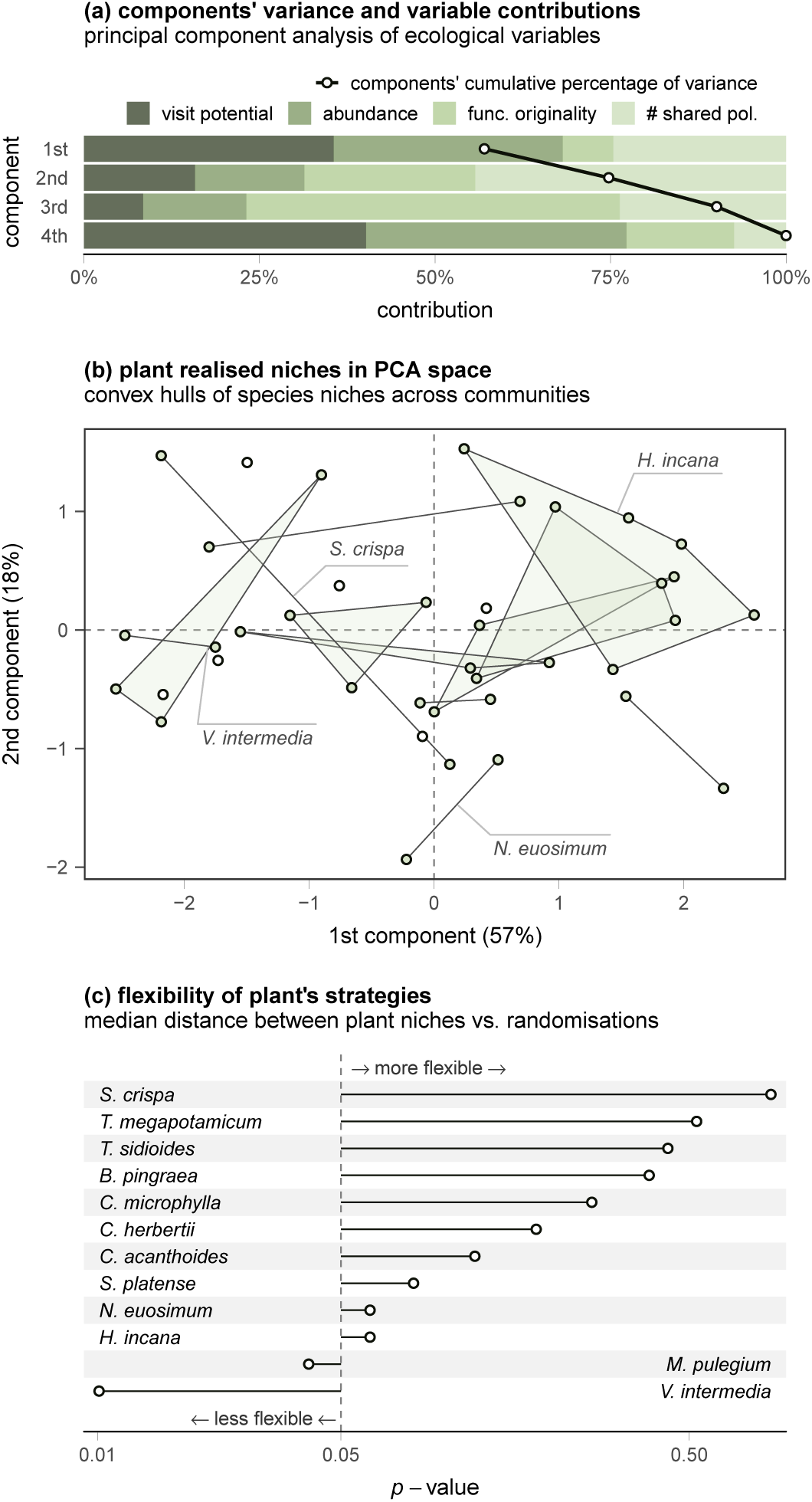
The flexibility of plant strategies. (a) The two first components explain a large proportion of the total variance. (b) When plants that were sampled in more than one community are plotted in terms of these two components, we observe that their points—which represent the strategy (the particular combination of ecological factors) of that species in its community—do not seem to be grouped by plant species. (c) This was confirmed using Monte Carlo randomizations of the median distance between strategies of a plant species. Only two of the examined species had strategies that were less flexible than would be expected at random.

## Discussion

Our results suggest that community context plays a central role in determining the pollen deposition dynamics and ultimately the net cost or benefit of sharing pollinators. First, we found that multiple ecological factors can modulate the quality of the pollination service; however, conspecific and heterospecific pollen deposition are tightly coupled and this creates a clear trade-off between the quantity and purity of pollination (Thomson et al. 2019). Second, we found that the way these factors shape pollen deposition for a species could be dramatically different across communities. For instance, while a plant species in a particular community could show high levels of pollinator sharing and relatively low trait differentiation, the same species in another community can have relatively high trait differentiation and low levels of pollinator sharing. Our findings highlight that trade-offs can at least partially explain the coexistence of facilitative and competitive effects of animal-mediated pollination in the pollination service.

The trade-offs involved in attaining high-quality pollination service (and more broadly between facilitation and competition) are likely to arise when plants simultaneously maximise the deposition of conspecific pollen and minimise that of heterospecific pollen. In the short term, being a specialist and sharing no pollinators might reduce competition (Muchhala et al. 2010) and hence be preferable. This may be due to both costs to male fitness (Morales and Traveset 2008, Muchhala and Thomson 2012), and also, as we show here, because sharing pollinators reduces both the quantity and purity of the conspecific pollen deposited. However, over long periods of time, there could be a risk associated with a specialist plant having few pollinators (Ricketts 2004). To ensure long-term survival, it is thus likely that plants also need to balance this risk with the costs of sharing pollinators (Aizen et al. 2012). One possible solution is to share pollinators *and* have original traits—as we show that trait originality is generally beneficial to pollen deposition and it is commonly thought that species that are further from others in trait space benefit from reduced competition. Yet, there are two possible caveats to this strategy that highlight the interrelatedness of the ecological factors. First, in a mutualism context, it is also possible that trait originality could come at the cost of being less ‘apparent’ to pollinators (Reverté et al. 2016). Second, the negative relationship between originality and generalism (Carvalheiro et al. 2014) has been shown to depend on plant abundance (Coux et al. 2016), with generalist species being able to have original traits only when they are also abundant enough to provide a valuable reward to make visiting worthwhile to pollinators.

Visit potential (high pollen and visits) and floral abundance, which were the most important predictors of pollen deposition here, introduced an even more explicit trade-off between gaining conspecific pollen and avoiding heterospecific pollen. Receiving high visitation increases conspecific pollen deposition but increases heterospecific pollen deposition to a greater extent—even when the visitors are likely to carry a high proportion of conspecific pollen (Fang and Huang 2016). Contrastingly, being abundant reduces the amount of conspecific pollen deposited and simultaneously reduces heterospecific pollen at a faster rate. Our results corroborate the importance that two-species studies have ascribed to visitation and abundance (Feldman et al. 2004, Muñoz and Cavieres 2008, Morales and Traveset 2008), but they also suggest that (because visitation, pollen production and abundance are usually correlated; Sargent and Otto 2006) balancing the pros and cons of sharing pollinators at the community level is not trivial. The fact that no species can easily outcompete others for pollination might be partially responsible for the diversity of plant-pollinator communities (Benadi and Pauw 2018).

We observed, as expected, that the effects of pollen deposition can vary widely among species. For instance, the fitness of some plant species can be hurt even by low amounts of heterospecific pollen, while the fitness of others can instead be limited by the amount of conspecific pollen (Campbell and Motten 1985, Arceo-Gómez et al. 2019). Alternatively, plant species can also differ substantially in the extent to which self- vs. outcross-pollen differ in their value for fertilization. The difference can be particularly relevant for species that are not self-fertile or those in which self-fertilization is rarely effective due to a temporary separation in the maturation of the sexes (dichogamy).

Importantly, we show here that the balances between costs and benefits are determined not only by species identity but also by the community to which plants belong. Specifically, most plant species appear to be flexible enough to adopt markedly different “strategies” in different communities. From an evolutionary perspective, our results suggest that selection for a particular strategy might say something about the community in which a species has typically inhabited during its evolutionary history. Furthermore, from a more applied perspective, flowering plants are sometimes introduced to attract pollinators on other nearby plants. On the one hand, our results suggest that plants that increase the relative originality of natives (e.g. through distinct phenology) might have positive effects (Gibson et al. 2012). On the other, because different strategies can lead to different outcomes across communities, our results also highlight the difficulties involved in predicting whether the introduced plant species will facilitate or compete with neighbours (Bartomeus et al. 2008). Other factors that we were unable to measure (e.g. pollinator behaviour and densities or the spatial context) have also been shown to play a role in the outcome of animal-mediated pollination (Cariveau and Norton 2009, Flanagan et al. 2011, Ye et al. 2014, Thomson et al. 2019). Nevertheless, our results indicate that the strategies a plant might use to successfully minimise competition for pollination (or maximise facilitation) must be determined relative to other species in the community, rather than an absolute property of the species itself.

Overall, using empirical data on pollen deposition, we show at the community level that sharing pollinators has a smaller effect on pollen deposition than what we expected based on experimental studies with a handful species. Other factors that underpin community dynamics (abundance, traits, visitation) also influence patterns of pollination quantity and purity. The interrelatedness of these factors, and the flexibility of species to position themselves across them, means that their contributions to the quality of the pollination service cannot be understood in isolation. All of the factors we analysed involve substantial trade-offs in pollen deposition in the short and likely also in the long term. These trade-offs emphasise the inherently competitive nature of pollination. However, many of the widely used theoretical models of plant-pollinator communities do not account for the adverse effects of sharing pollinators (but see Rohr et al. 2014 and similar). We therefore propose that achieving a better understanding of species coexistence and how pollination supports plant biodiversity will require seeing them as both mutualistic and competitive communities (Johnson and Bronstein 2019).

## Acknowledgements

We thank Jamie Stavert, Bernat Bramon Mora, Laís Maia, and Michelle Marraffini for feedback and valuable discussions. We also thank Cátedra de Botánica General, Facultad de Agronomía, Universidad de Buenos Aires, the Agrasar and Bordeu families, and the University of Buenos Aires, for logistical support and permission to conduct this study at estancias Anquilóo, Las Chilcas and San Claudio, respectively. Fieldwork was supported by grants PICT 08–12504 and 0851. EFC acknowledges the support from the University of Canterbury Doctoral Scholarship and a New Zealand International Doctoral Research Scholarship administered by Education New Zealand. DBS and JMT acknowledge the support of Rutherford Discovery Fellowships (RDF-13-UOC-003 and RDF-UOC-1002) and the Marsden Fund Council (UOC-1705), administered by the Royal Society of New Zealand Te Apārangi.

## Notes

https://github.com/efcaguab/pollen-competition

## References

Aizen, M. A., and A. E. Rovere. 2010. Reproductive interactions mediated by flowering overlap in a temperate hummingbird-plant assemblage. Oikos 119:696–706.

Aizen, M. A., M. Sabatino, and J. M. Tylianakis. 2012. Specialization and rarity predict nonrandom loss of interactions from mutualist networks. Science 335:1486–1489.

Arceo-Gómez, G., and T.-L. Ashman. 2016. Invasion status and phylogenetic relatedness predict cost of heterospecific pollen receipt: Implications for native biodiversity decline. Journal of Ecology 104:1003–1008.

Arceo-Gómez, G., R. L. Kaczorowski, C. Patel, and T.-L. Ashman. 2019. Interactive effects between donor and recipient species mediate fitness costs of heterospecific pollen receipt in a co-flowering community. Oecologia 189:1041–1047.

Ashman, T.-L., and G. Arceo-Gómez. 2013. Toward a predictive understanding of the fitness costs of heterospecific pollen receipt and its importance in co-flowering communities. American Journal of Botany 100:1061–1070.

Bartomeus, I., M. Vilà, and L. Santamaría. 2008. Contrasting effects of invasive plants in plant-pollinator networks. Oecologia 155:761–770.

Bascompte, J., and P. Jordano. 2007. Plant-Animal Mutualistic Networks: The Architecture of Biodiversity. Annual Review of Ecology, Evolution, and Systematics 38:567–593.

Bascompte, J., P. Jordano, and J. M. Olesen. 2006. Asymmetric Coevolutionary Networks Facilitate Biodiversity Maintenance. Science 312:431–433.

Benadi, G., and A. Pauw. 2018. Frequency dependence of pollinator visitation rates suggests that pollination niches can allow plant species coexistence. Journal of Ecology 106:1892–1901.

Bosch, J., A. M. Martín González, A. Rodrigo, and D. Navarro. 2009. Plant-pollinator networks: Adding the pollinator’s perspective. Ecology Letters 12:409–419.

Campbell, D. R., and A. F. Motten. 1985. The Mechanism of Competition for Pollination between Two Forest Herbs. Ecology 66:554–563.

Cariveau, D. P., and A. P. Norton. 2009. Spatially contingent interactions between an exotic and native plant mediated through flower visitors. Oikos 118:107–114.

Caruso, C. M. 2000. Competition for Pollination Influences Selection on Floral Traits of Ipomopsis aggregata. Evolution 54:1546–1557.

Carvalheiro, L. G., J. C. Biesmeijer, G. Benadi, J. Fründ, M. Stang, I. Bartomeus, C. N. Kaiser-Bunbury, M. Baude, S. I. F. Gomes, V. Merckx, K. C. R. Baldock, A. T. D. Bennett, R. Boada, R. Bommarco, R. Cartar, N. Chacoff, J. Dänhardt, L. V. Dicks, C. F. Dormann, J. Ekroos, K. S. Henson, A. Holzschuh, R. R. Junker, M. Lopezaraiza-Mikel, J. Memmott, A. Montero-Castaño, I. L. Nelson, T. Petanidou, E. F. Power, M. Rundlöf, H. G. Smith, J. C. Stout, K. Temitope, T. Tscharntke, T. Tscheulin, M. Vilà, and W. E. Kunin. 2014. The potential for indirect effects between co-flowering plants via shared pollinators depends on resource abundance, accessibility and relatedness. Ecology Letters 17:1389–1399.

Coux, C., R. Rader, I. Bartomeus, and J. M. Tylianakis. 2016. Linking species functional roles to their network roles. Ecology Letters 19:762–770.

Dafni, A., and D. Firmage. 2000. Pollen viability and longevity: Practical, ecological and evolutionary implications. Plant Systematics and Evolution 222:113–132.

Fang, Q., and S.-Q. Huang. 2016. A paradoxical mismatch between interspecific pollinator moves and heterospecific pollen receipt in a natural community. Ecology 97:1970–1978.

Feinsinger, P. 1987. Effects of plant species on each others pollination: Is community structure influenced? Trends in Ecology & Evolution 2:123–126.

Feldman, T. S., W. F. Morris, and W. G. Wilson. 2004. When can two plant species facilitate each other’s pollination? Oikos 105:197–207.

Flanagan, R. J., R. J. Mitchell, and J. D. Karron. 2011. Effects of multiple competitors for pollination on bumblebee foraging patterns and Mimulus Ringens reproductive success. Oikos 120:200–207.

Gibson, M. R., D. M. Richardson, and A. Pauw. 2012. Can floral traits predict an invasive plant’s impact on native plant-pollinator communities? Journal of Ecology 100:1216–1223.

Johnson, C. A., and J. L. Bronstein. 2019. Coexistence and competitive exclusion in mutualism. Ecology 100:e02708.

de Jong, T. J., N. M. Waser, M. V. Price, and R. M. Ring. 1992. Plant size, geitonogamy and seed set in Ipomopsis aggregata. Oecologia 89:310–315.

Klein, A.-M., B. E. Vaissiere, J. H. Cane, I. Steffan-Dewenter, S. A. Cunningham, C. Kremen, and T. Tscharntke. 2007. Importance of pollinators in changing landscapes for world crops. Proceedings of the Royal Society B: Biological Sciences 274:303–313.

Laliberté, E., and P. Legendre. 2010. A distance-based framework for measuring functional diversity from multiple traits. Ecology 91:299–305.

Laliberté, E., P. Legendre, and B. Shipley. 2014. FD: Measuring functional diversity from multiple traits, and other tools for functional ecology.

Marrero, H. J., D. Medan, G. Zarlavsky, and J. P. Torretta. 2016. Agricultural land management negatively affects pollination service in Pampean agro-ecosystems. Agriculture, Ecosystems & Environment 218:28–32.

Marrero, H. J., J. P. Torretta, and D. Medan. 2014. Effect of land use intensification on specialization in plant-floral visitor interaction networks in the Pampas of Argentina. Agriculture, Ecosystems & Environment 188:63–71.

Marrero, H. J., J. P. Torretta, D. P. Vázquez, K. Hodara, and D. Medan. 2017. Exotic plants promote pollination niche overlap in an agroecosystem. Agriculture, Ecosystems & Environment 239:304–309.

Memmott, J. 1999. The structure of a plant-pollinator food web. Ecology Letters 2:276–280.

Morales, C. L., and A. Traveset. 2008. Interspecific pollen transfer: Magnitude, prevalence and consequences for plant fitness. Critical Reviews in Plant Sciences 27:221–238.

Muchhala, N., and J. D. Thomson. 2012. Interspecific competition in pollination systems: Costs to male fitness via pollen misplacement: Pollen misplacement. Functional Ecology 26:476–482.

Muchhala, N., Z. Brown, W. S. Armbruster, and M. D. Potts. 2010. Competition Drives Specialization in Pollination Systems through Costs to Male Fitness. The American Naturalist 176:732–743.

Muñoz, A. A., and L. A. Cavieres. 2008. The presence of a showy invasive plant disrupts pollinator service and reproductive output in native alpine species only at high densities: Invasive impacts on native species pollination. Journal of Ecology 96:459–467.

Ollerton, J., R. Winfree, and S. Tarrant. 2011. How many flowering plants are pollinated by animals? Oikos 120:321–326.

Pinheiro, J., D. Bates, S. DebRoy, D. Sarkar, and R Core Team. 2018. Nlme: Linear and Nonlinear Mixed Effects Models.

Ramsey, M., and G. Vaughton. 2000. Pollen quality limits seed set in Burchardia Umbellata (Colchicaceae). American Journal of Botany 87:845–852.

Reverté, S., J. Retana, J. M. Gómez, and J. Bosch. 2016. Pollinators show flower colour preferences but flowers with similar colours do not attract similar pollinators. Annals of Botany 118:249–257.

Ricketts, T. H. 2004. Tropical Forest Fragments Enhance Pollinator Activity in Nearby Coffee Crops. Conservation Biology 18:1262–1271.

Rohr, R. P., S. Saavedra, and J. Bascompte. 2014. On the structural stability of mutualistic systems. Science 345:1253497.

Sargent, R. D., and S. P. Otto. 2006. The role of local species abundance in the evolution of pollinator attraction in flowering plants. The American Naturalist 167:67–80.

Seifan, M., E.-M. Hoch, S. Hanoteaux, and K. Tielbörger. 2014. The outcome of shared pollination services is affected by the density and spatial pattern of an attractive neighbour. Journal of Ecology 102:953–962.

Stavert, J. R., I. Bartomeus, J. R. Beggs, A. C. Gaskett, and D. E. Pattemore. 2019. Plant species dominance increases pollination complementarity and plant reproductive function. Ecology 100.

Thébault, E., and C. Fontaine. 2008. Does asymmetric specialization differ between mutualistic and trophic networks? Oikos 117:555–563.

Thomson, J. D., H. F. Fung, and J. E. Ogilvie. 2019. Effects of spatial patterning of co-flowering plant species on pollination quantity and purity. Annals of Botany 123:303–310.

Tur, C., A. Sáez, A. Traveset, and M. A. Aizen. 2016. Evaluating the effects of pollinatormediated interactions using pollen transfer networks: Evidence of widespread facilitation in south Andean plant communities. Ecology Letters 19:576–586.

Vázquez, D. P., C. J. Melián, N. M. Williams, N. Blüthgen, B. R. Krasnov, and R. Poulin. 2007. Species Abundance and Asymmetric Interaction Strength in Ecological Networks. Oikos 116:1120–1127.

Waser, N. M. 1978. Interspecific pollen transfer and competition between co-occurring plant species. Oecologia 36:223–236.

Yang, S., M. J. Ferrari, and K. Shea. 2011. Pollinator behavior mediates negative interactions between two congeneric invasive plant species. The American Naturalist 177:110–118.

Ye, Z.-M., W.-K. Dai, X.-F. Jin, R. W. Gituru, Q.-F. Wang, and C.-F. Yang. 2014. Competition and facilitation among plants for pollination: Can pollinator abundance shift the plant-plant interactions? Plant Ecology 215:3–13.

