## Supplementary Information for "The trade-offs of sharing pollinators: pollination service is determined by the community context"

*E. Fernando Cagua, Hugo J. Marrero, Jason M. Tylianakis, Daniel B. Stouffer*

Table S1: Summary of the model used to analyse the relationship between heterospecific and conspecific pollen

| predictor | estimate | S.E. | z-value |
|---|---|---|---|
| **fixed component** | | | |
| (Intercept) | 4.976 | 0.279 | 17.862 |
| heterospecific | 0.008 | 0.017 | 0.474 |
| **random component (species:community)** | | | |
| S.D. random intercept | 1.964 | - | - |
| S.D. random slope | 0.120 | - | - |

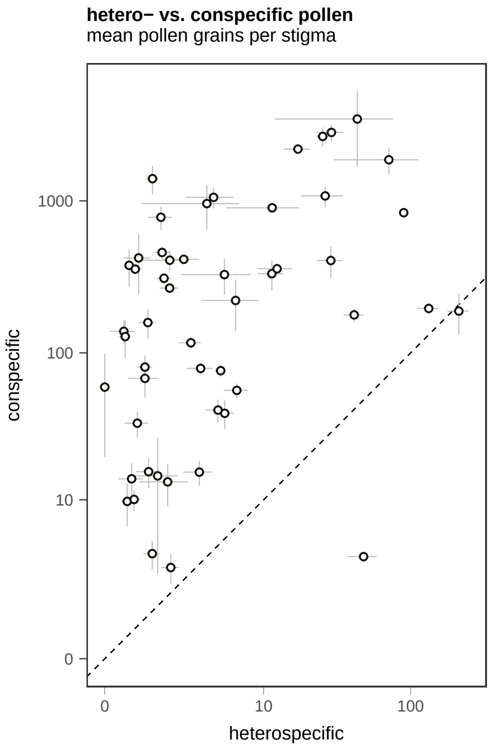


Figure S1: Despite the variation in these slopes, plants overall had more conspecific than heterospecific pollen deposited in their stigmas.

Table S2: The slope of the relationship between heterospecific and conspecific pollen for each species in their community (fixed effect + conditional effect). Community names are constructed by location - agricultural/restored - fragment number.

| species name | community | slope | S.E. |
|---|---|---|---|
| *Aloysia gratissima* | Anquilóo - reserve - 2 | 0.0746 | 0.0144 |
| *Baccharis pingraea* | San Claudio - reserve - 1 | -0.0012 | 0.0359 |
| *Carduus acanthoides* | Anquilóo - agricultural - 2 | 0.0116 | 0.0147 |
| *Carduus acanthoides* | San Claudio - agricultural - 1 | -0.0106 | 0.0040 |
| *Carduus acanthoides* | San Claudio - agricultural - 2 | 0.0518 | 0.0044 |
| *Carduus acanthoides* | San Claudio - reserve - 1 | 0.0781 | 0.0710 |
| *Carduus acanthoides* | San Claudio - reserve - 2 | -0.0008 | 0.0359 |
| *Cirsium vulgare* | Anquilóo - agricultural - 2 | -0.0401 | 0.0025 |
| *Cirsium vulgare* | Las Chilcas - reserve - 1 | 0.0007 | 0.0012 |
| *Cirsium vulgare* | San Claudio - agricultural - 2 | 0.0197 | 0.0158 |
| *Cirsium vulgare* | San Claudio - reserve - 1 | -0.0149 | 0.0076 |
| *Condalia microphylla* | Anquilóo - reserve - 1 | 0.0487 | 0.0200 |
| *Cypella herbertii* | Las Chilcas - agricultural - 2 | 0.0037 | 0.0002 |
| *Cypella herbertii* | Las Chilcas - reserve - 1 | -0.0052 | 0.0001 |
| *Descurania argentina* | Anquilóo - agricultural - 2 | 0.0429 | 0.0048 |
| *Diplotaxis tenuifolia* | Anquilóo - reserve - 1 | 0.0008 | 0.0004 |
| *Diplotaxis tenuifolia* | Anquilóo - reserve - 2 | 0.5173 | 0.0270 |
| *Diplotaxis tenuifolia* | San Claudio - reserve - 2 | -0.0045 | 0.0001 |
| *Dipsacus sp.* | San Claudio - reserve - 2 | -0.0368 | 0.0648 |
| *Gaillardia megapotamica* | Anquilóo - reserve - 2 | 0.0016 | 0.0004 |
| *Glandularia hookeriana* | Anquilóo - reserve - 2 | -0.0942 | 0.0244 |
| *Hirschfeldia incana* | Anquilóo - agricultural - 1 | -0.0045 | 0.0013 |
| *Hirschfeldia incana* | Anquilóo - agricultural - 2 | -0.0148 | 0.0057 |
| *Hirschfeldia incana* | San Claudio - agricultural - 1 | 0.0110 | 0.0020 |
| *Hirschfeldia incana* | San Claudio - agricultural - 2 | 0.0031 | 0.0023 |
| *Hirschfeldia incana* | San Claudio - reserve - 1 | 0.0022 | 0.0002 |
| *Hirschfeldia incana* | San Claudio - reserve - 2 | 0.0432 | 0.0020 |
| *Lycium chilense* | Anquilóo - reserve - 2 | -0.3355 | 0.0087 |
| *Mentha pulegium* | Las Chilcas - agricultural - 2 | 0.0136 | 0.0866 |
| *Mentha pulegium* | Las Chilcas - reserve - 1 | 0.3973 | 0.0388 |
| *Nierembergia aristata* | Anquilóo - agricultural - 1 | 0.0197 | 0.0217 |
| *Nierembergia aristata* | Anquilóo - reserve - 1 | -0.0065 | 0.0016 |
| *Nierembergia aristata* | Anquilóo - reserve - 2 | -0.0048 | 0.0011 |
| *Nothoscordum euosimum* | Las Chilcas - agricultural - 1 | 0.0405 | 0.0034 |
| *Nothoscordum euosimum* | Las Chilcas - agricultural - 2 | -0.0045 | 0.1162 |
| *Physalis viscosa* | Anquilóo - agricultural - 1 | 0.0041 | 0.0005 |
| *Prosopidastrum globosum* | Anquilóo - reserve - 2 | -0.0012 | 0.0194 |
| *Senecio pulcher* | Las Chilcas - agricultural - 1 | -0.0104 | 0.0007 |
| *Sisyrinchium platense* | Las Chilcas - agricultural - 1 | -0.2850 | 0.0203 |
| *Sisyrinchium platense* | Las Chilcas - agricultural - 2 | -0.0487 | 0.0324 |
| *Sisyrinchium platense* | Las Chilcas - reserve - 1 | 0.0206 | 0.1143 |
| *Solanum sisymbriifolium* | San Claudio - agricultural - 1 | 0.0002 | 0.0004 |
| *Sphaeralcea crispa* | Anquilóo - reserve - 1 | -0.0601 | 0.0133 |
| *Stemodia lanceolata* | Las Chilcas - agricultural - 1 | -0.0044 | 0.0001 |
| *Thelesperma megapotamicum* | Anquilóo - agricultural - 1 | -0.0022 | 0.0025 |
| *Turnera sidioides* | Anquilóo - agricultural - 1 | -0.0002 | 0.0001 |
| *Turnera sidioides* | Anquilóo - agricultural - 2 | -0.0140 | 0.0170 |
| *Turnera sidioides* | Anquilóo - reserve - 2 | -0.0014 | 0.0002 |
| *Verbena intermedia* | Anquilóo - reserve - 2 | -0.0643 | 0.0327 |
| *Verbena intermedia* | San Claudio - agricultural - 2 | 0.0932 | 0.0071 |
| *Verbena intermedia* | San Claudio - reserve - 2 | -0.0073 | 0.0101 |

Table S3: The coefficient of determination $R^2$ of the most parsimonious pollen deposition models (those with the lowest AICc). The marginal coefficient of determination describes the proportion of variance explained by just the fixed effects.

| conditional $R^2_{(c)}$ | | | marginal $R^2_{(m)}$ | | |
|---|---|---|---|---|---|
| mean | min | max | mean | min | max |
| **conspecific pollen** | | | | | |
| 0.91 | 0.87 | 0.93 | 0.09 | 0.06 | 0.14 |
| **heterospecific pollen** | | | | | |
| 0.80 | 0.76 | 0.87 | 0.27 | 0.21 | 0.35 |

Table S4: Comparison of the two random structures we considered for the models of conspecific and heterospecific pollen deposition. The table shows median ΔAIC values of 99 bootstrap resamples of the data. The 5th and 95th percentile are shown inside square brackets. Communities are defined by individual fragments but ignore the hierarchical arrangement of sampling sites.

| | ΔAIC | |
|---|---|---|
| random structure | median | C.I. |
| **conspecific pollen** | | |
| 1 \| plant sp. * community | 0.0 | [0, 0] |
| 1 \| plant sp. | 30.7 | [8.2, 58.1] |
| **heterospecific pollen** | | |
| 1 \| plant sp. * community | 0.0 | [0, 0] |
| 1 \| plant sp. | 44.6 | [19.3, 88.4] |

Table S5: Comparison of the different fixed structures we considered for the models of conspecific and heterospecific pollen deposition. The table shows median ΔAIC values of 99 bootstrap resamples of the data. The 5th and 95th percentile are shown inside square brackets.

| | ΔAIC | |
|---|---|---|
| fixed structure | median | C.I. |
| **conspecific pollen** | | |
| ~ abundance + visit potential | 0.0 | [0, 0] |
| ~ abundance + visit potential + func. originality | 0.9 | [0.4, 1.3] |
| ~ abundance + visit potential + # shared pol. | 1.9 | [1.6, 2.1] |
| ~ abundance + visit potential + # shared pol. + func. originality | 2.2 | [1.6, 2.8] |
| ~ visit potential + func. originality | 2.8 | [2.1, 3.8] |
| ~ visit potential + # shared pol. + func. originality | 3.6 | [2.3, 4.6] |
| ~ visit potential | 118.3 | [75.3, 178.7] |
| ~ visit potential + # shared pol. | 119.0 | [76, 179.9] |
| ~ abundance | 189.7 | [150.1, 239.7] |
| ~ abundance + func. originality | 191.6 | [151.7, 241.6] |
| ~ abundance + # shared pol. | 191.7 | [151.9, 241.7] |
| ~ func. originality | 192.5 | [152.9, 242.2] |
| ~ abundance + # shared pol. + func. originality | 193.7 | [153.6, 243.6] |
| ~ # shared pol. + func. originality | 193.7 | [154.6, 243.7] |
| ~ # shared pol. | 351.8 | [293.5, 419.9] |
| **heterospecific pollen** | | |
| ~ abundance + visit potential | 0.0 | [0, 0] |
| ~ abundance + visit potential + func. originality | 1.1 | [0.5, 1.5] |
| ~ abundance + visit potential + # shared pol. | 2.1 | [1.9, 2.1] |
| ~ abundance + visit potential + # shared pol. + func. originality | 3.1 | [2.6, 3.5] |
| ~ visit potential + func. originality | 11.9 | [10, 13.9] |
| ~ visit potential + # shared pol. + func. originality | 13.2 | [11.2, 15.2] |
| ~ visit potential | 67.5 | [53.4, 87.5] |
| ~ visit potential + # shared pol. | 68.4 | [54.2, 88.7] |
| ~ abundance + # shared pol. | 206.9 | [160.6, 251.5] |
| ~ abundance | 207.6 | [162.8, 251.7] |
| ~ abundance + func. originality | 208.6 | [163.2, 252.6] |
| ~ abundance + # shared pol. + func. originality | 208.6 | [162.2, 253.2] |
| ~ func. originality | 214.3 | [168.3, 258.7] |
| ~ # shared pol. + func. originality | 216.3 | [170.3, 260.6] |
| ~ # shared pol. | 336.0 | [282.6, 391.5] |

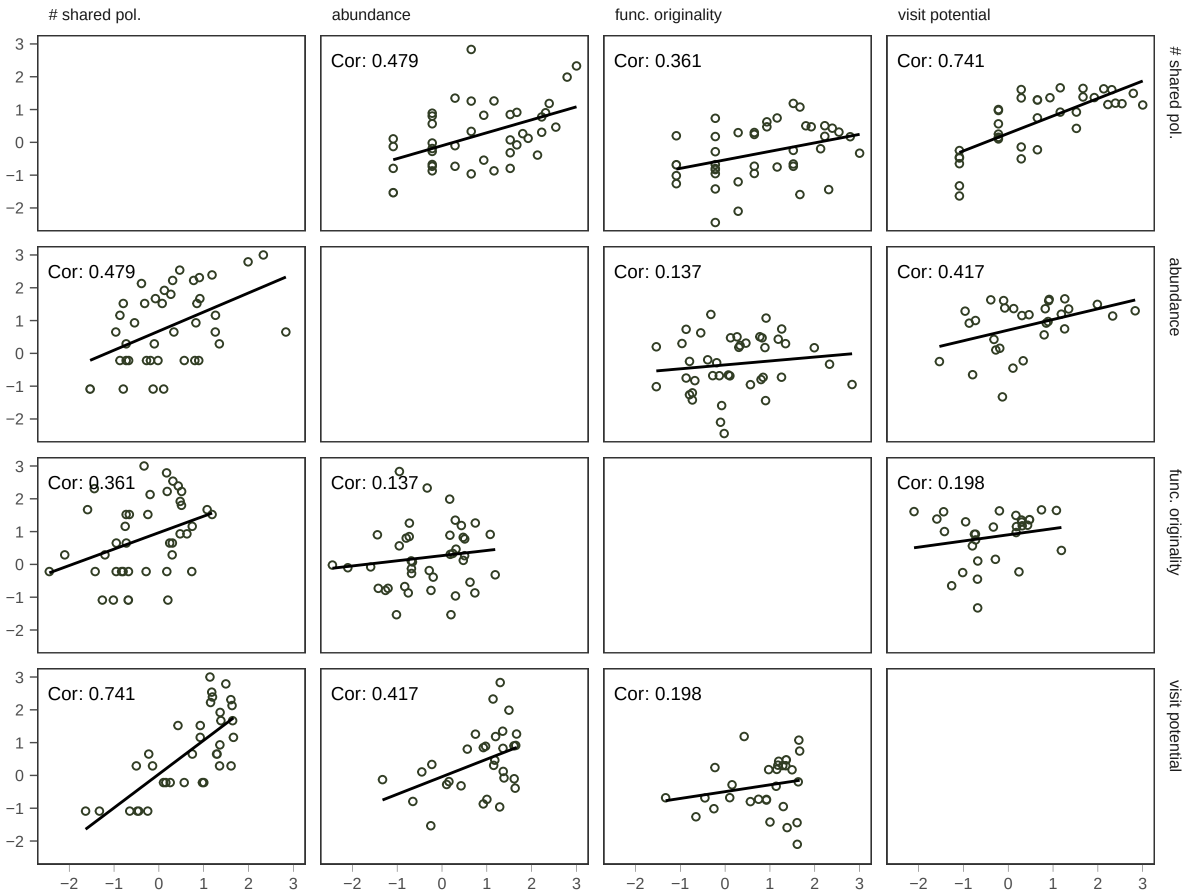


Figure S2: Correlation between the explanatory variables included in the statistical models.

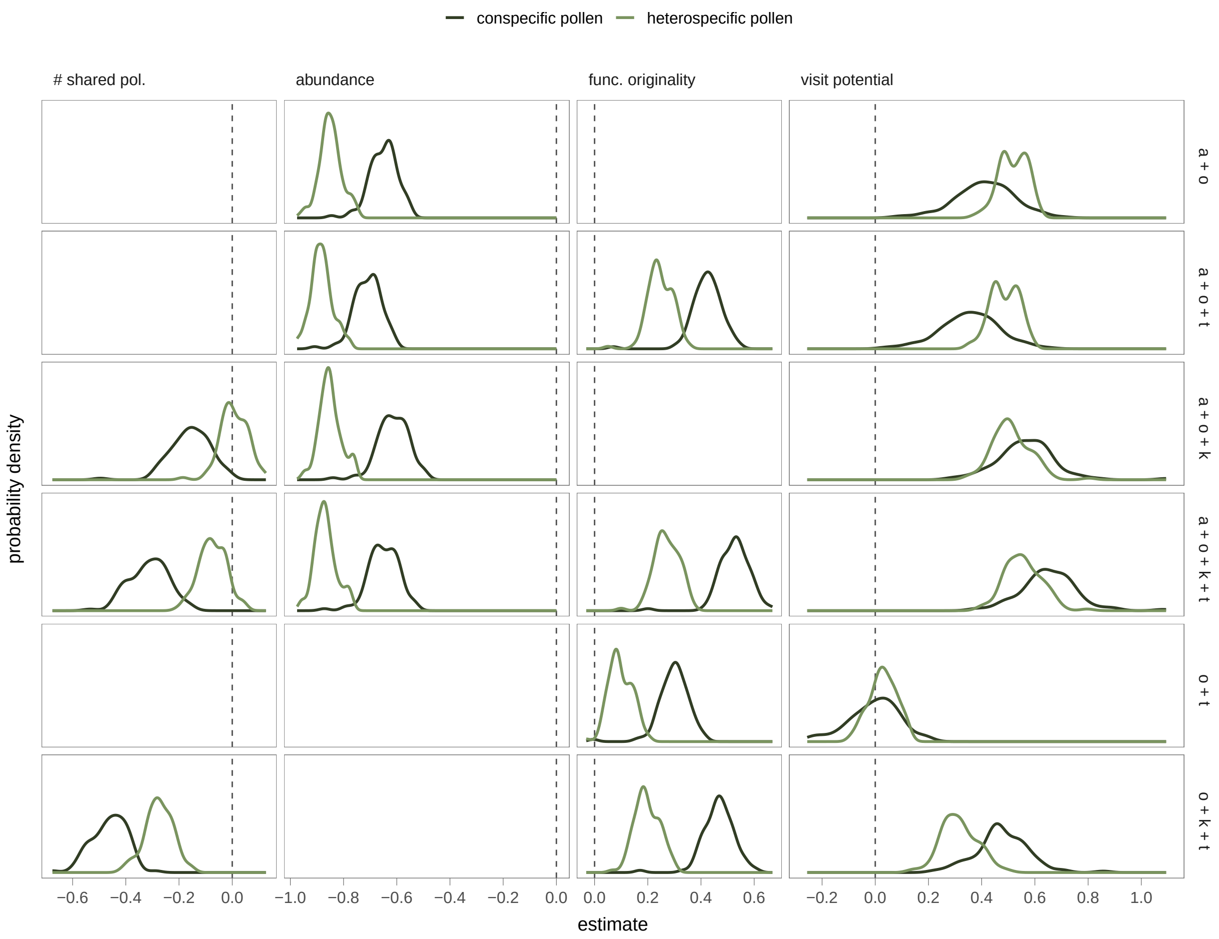


Figure S3: Distribution of effect estimates for models of conspecific and heterospecific pollen density gain. Model formulas have been abbreviated: $a$ for abundance, $k$ for the number of shared pollinators, $o$ for the visit potential, and $t$ for functional originality. Only candidate formulas with a $\Delta AICc < 4$ for either conspecific or heterospecific pollen are shown. Model candidates are arranged in decreasing order of support. Although relative abundance, the number of shared pollinators, and the visit potential were all positively correlated, the effect each had on conspecific pollen was similar among models that included all or just some of these three explanatory variables. One exception was visit potential, which exhibits a positive association with the relative amount of conspecific pollen under some variable combinations. Nevertheless, these differences were observed only in model specifications with relatively low AICc support.
